# The hummingbird and the hawk-moth: Species distribution, geographical partitioning, and macrocompetition across the United States

**DOI:** 10.1101/212894

**Authors:** Abdel Halloway, Christopher J. Whelan, Joel S. Brown

## Abstract

We introduce a new concept called macrocompetition – defined as the mutual suppression of diversity/species richness of competing clades – and investigate evidence for its existence. To this end, we analyzed the distribution of two convergent nectarivorous families, hawk-moths and hummingbirds, over the continental United States to determine whether there is geographic partitioning between the families and its potential causes. Using stepwise regression, we tested for latitudinal and longitudinal biases in the species richness of both taxa and the potential role of 10 environmental variables in their distribution pattern. Hawk-moth species richness increases with longitude (eastward-bias) while that of hummingbirds declines (westward-bias). Similar geographic patterns can be seen across Canada, Mexico and South America. Hawk-moth species richness is positively correlated with higher overall temperatures (especially summer minimums), atmospheric pressure, and summer precipitation; hummingbird species richness is negatively correlated with atmospheric pressure and positively correlated with winter daily maxima. The species richness patterns reflect each family’s respective anatomical differences and support the concept of macrocompetition between the two taxa. Hawk-moth species richness was highest in states with low elevation, summer-time flowering, and warm summer nights; hummingbird species richness is highest in the southwest with higher elevation, greater cool season flowering and high daytime winter temperatures. Hawk-moths and hummingbirds as distinct evolutionary technologies exhibit niche overlap and geographical partitioning. These are two of three indicators suggested by Brown and Davidson for inter-taxonomic competition. We intend the patterns revealed here to inspire further exploration into competition and community structuring between hawk-moths and hummingbirds.

## INTRODUCTION

Competitive interactions help shape distribution (Hutchinson 1978), origination (Rosenzweig 1978; Hutchinson 1978; Schluter 2000; Ripa et al. 2009) and extinction of species (Gause 1934). Competition affects small-scale interactions among species yet also drives larger scale phenomena and lies at the core of processes like competitive speciation (Rosenzweig 1978) and incumbent replacement (Rosenzweig and McCord, 1991; Silvestro et al., 2015). It is most often studied at the local scale, either between individuals within a population mutually suppressing fitness, or between populations mutually suppressing each other’s population size.

Competition may also operate at higher taxonomic levels. By occupying potential niches space of another, one taxonomic group may limit the species diversification or adaptive radiation of another. In this case, competition suppresses species richness rather than fitness or population size. We propose that competition thus acts on three levels:

- Microcompetition operates between individuals and suppresses access to resources
- Mesocompetition operates between populations and suppresses population sizes
- Macrocompetition operates between higher order taxa and suppresses species richness

These three forms of competition should occur on different temporal, spatial and taxonomic scales. Macrocompetition, which suppresses species diversity and the radiation of species within taxonomic groups, must occur over large temporal and spatial scales and at taxonomic levels higher than the species. Because of this link between spatial, temporal, and organizational scales, macrocompetition must be studied at its own appropriate scale (Jablonski, 2008). Just as population level mesocompetition is not studied by aggregating individual microcompetitive interactions, macrocompetition cannot be studied through the aggregation of mesocompetitive and microcompetitive interactions.

When studying macrocompetition, the adaptations specific to each clade are most important. These clade-specific adaptations, which we refer to as evolutionary technologies, are the tools that allow each clade to exploit environments and resources and form the basis of macrocompetition. For macrocompetition, each taxa must exhibit one or more derived traits that are shared among the members of the taxa but distinct from members of the competing taxa. The evolutionary feasibility of these traits to the members of the taxa; and their unavailability to members of other taxa defines the evolutionary technology (Vincent and Brown 2005). While not originally intended as such, Families may represent a rough, but good first cutoff for describing different evolutionary technologies (Pintor et al. 2011); and certainly members of different Orders, Classes and Phyla represent different taxa for the purposes of macrocompetition.

Mesocompetition between populations of different taxa has been well-documented. Examples include tadpoles and aquatic insects (Morin et al., 1988) and insect larvae (Mokany and Shine, 2003), granivorous rodents and ants (Brown and Davidson, 1977; Brown and Davidson, 1979), granivorous birds and rodents (Brown et al., 1997), frugivorous birds and bats (Palmeirim et al., 1989), insectivorous lizards and birds (Wright, 1979), and insectivorous birds and ants (Haeming, 1994; Jedlicka et al. 2006). Mesocompetition may even exist between species of separate phyla, such as the competition between scavenging vertebrates and microbes for detritus (Janzen, 1977; Shivik 2006) or vertebrates and fungi for rotting fruit (Cipollini and Stiles 1993; Cipollini and Levey 1997). Brown and Davidson (1977) identified three key indicators to determine potential intertaxonomic mesocompetition: 1) shared extensive use of the same particular resource, 2) reciprocal increases in population size when one competing species is excluded, and 3) partitioning along a geographic or climatic gradient. We propose analogous indicators as signals of macrocompetition: 1) shared extensive use of the same class of resources, 2) reciprocal increases in species richness via adaptive radiation when a competing taxon is excluded, and 3) partitioning along geographical and climatic gradients across the shared taxa’s range.

Pollination systems provide ample opportunities for intertaxonomic competition. Both Primack and Howe (1975) and Thomas et al. (1986) reported competition between hummingbirds and butterflies, and Laverty and Plowright (1985) reported competition between hummingbirds and bumblebees. Due to many convergent characteristics, competition between hawk-moths (Sphingidae) and hummingbirds (Trochilidae) seem just as likely. Both taxa are highly-specialized nectar feeders and pollinators as adults. They have similar sizes, hover when feeding, and some species in each taxon possess tongues and other features that are often adapted to a single species of plant (Johnsgard, 1997; Tuttle, 2007). Despite their remarkable similarity and strong niche overlap, competition between these two Families has seldom been investigated. Only Carpenter (1979) explored the possibility of direct competition between hawk-moths and hummingbirds. Her study documented spatial and temporal partitioning between hawk-moths and hummingbirds. Hawk-moths dominated *Ipomopsis* feeding sites through depletion of nectar resources. Hummingbirds exhibited aggressive behaviour towards hawk-moths, suggesting hummingbirds perceive hawk-moths as competitors.

Differences in morphology and physiology can shape broad scale biogeographical patterns (Buckley et al., 2012). With this in mind, we compared species richness of hummingbirds and hawk-moths at the continental scale of North America with the goal of evaluating hallmarks of macrocompetition as modified from Brown and Davidson (1977). We seek broad scale geographic and climatic correlations of diversity that might provide insights into the patterns of diversity of hawk-moths and hummingbirds that may result from inter-taxon competition. Do diversity patterns of these two families covary positively or negatively? As nocturnal ectotherms, does hawk-moth diversity increase with summer rain and temperatures? As diurnal endotherms, do hummingbirds gain a competitive edge with colder temperature and cool season flowering? Do hawk-moths suffer more from low oxygen and elevation than do hummingbirds? Ultimately, to what extent can large-scale biogeography provide insights and clues into competition and geographical partitioning?

## MATERIALS AND METHODS

### Study Families

Hawk-moths (Order Lepidoptera, Family Sphingidae) and Hummingbirds (Order Apodiformes, Family Trochilidae) are nectarivores exhibiting morphological convergence. Worldwide, the approximately 953 species of hawk-moths (Kitching, and Cadiou, 2000) are moderate to large sized insects with wingspans that range from 25 to 200 mm (Kitching and Cadiou, 2000) and body weights ranging from 0.1 to 7 g (Janzen 1984). Hawk-moths outside of the tribe Smerinthini typically possess enhanced proboscides for nectar feeding and water drinking, allowing a longer lifespan than species which survive on fat reserves during their adult phase of the life cycle (Janzen, 1984). Locally, they seem to distinguish between visited and unvisited flowers. Over the course of days, they efficiently revisit flowers and patches on a regular basis and, over seasons, exhibit well directed long distance movements and migrations (Janzen, 1984). Hawk-moths evolved unique flight skills, including the ability to hover and a capacity for quick, long distance flight (Scoble, 1992). For instance, about half of the hawk-moth species at Santa Rosa National Park in Costa Rica migrate out of the park (Janzen, 1986). Many North America species disperse across continents, though the consistency and regularity of such dispersals are unknown (Tuttle, 2007). Some North American species likely migrate between North and South America as such cross-continental migration is known for many hawk-moths of the Western Palearctic (Pittaway, 1993).

All 328 hummingbird species reside in the New World (Schumann, 1999). The family includes the smallest known bird species with wing lengths from 29 to ≥ 90 mm (Johnsgard, 1997) and body masses ranging from 2 to 21 g (Schumann, 1999). Hummingbirds, like hawk-moths, possess specialized features for nectar-feeding, including elongated bills and extensible bitubular tongues for reaching and extracting nectar. Large breast muscles (30% of body weight) and specialized wings giving them the ability to hover and fly backwards. Hummingbirds are capable of long distance flight, with 13 of the 15 species of the United States exhibiting some degree of long distance migration (Johnsgard, 1997).

Many New World species of flowers exhibit distinct pollination syndromes that favor the morphology and behavior of hawk-moths or hummingbirds, respectively. Phalaenophilic (moth-pollinated) flowers typically open at night and use odor instead of visual cues as attractants, resulting in strongly scented but pale flowers. Comparatively narrow nectar tubes match the thin probosci typical of moths. Sex organs of phalaenophilic flowers are typically recessed, with the anther and stigma inserted within the corolla tube.

Ornithophilic (hummingbird-pollinated) flowers typically open during the day, are vividly colored (usually red), and offer little to no scent. Relatively wide nectar tubes match the bill width typical of hummingbirds (Faegri and van der Pijl, 1979). Exserted sex organs with the anther and stigma extending beyond the corolla characterize ornithophilous flowers (Kulbaba and Morley, 2008).

Despite these differences in flower morphology, both phalaenophilic and ornithophilic flowers share traits through convergent evolution. Both offer abundant nectar sources contained deep within long nectar tubes. Visual guides for pollinators are relatively absent in both flower types, with moths using the contours of the blossom as a guide (Faegri and van der Pijl, 1979). Individuals of both hawk-moths and hummingbirds prefer high sugar and abundant nectar, and each family readily feeds on the other’s flowers (Cruden et al., 1983; Cruden et al., 1976; Haber and Frankie, 1989). This extensive niche overlap offers ample opportunities for competition.

### Distribution Analysis

We determined the species richness of hummingbirds and hawk-moths across the continental USA. Using range maps and text descriptions provided by Johnsgard (1997) and Tuttle (2007), we determined the species richness for the 49 states of the continental USA. We used states as our scale of resolution as finer scale distribution data for hawk-moths does not exist. For our purpose of looking at large-scale biogeographic patterns, coarse but complete data is more important than fine but incomplete data. Our analysis included rare native species but excluded species non-native to the United States. First, we tested for geographic gradients of each Family’s diversity by creating a general linear model in which each state’s longitude and latitude – based upon the centroid of the state – were independent variables and species richness as the dependent variable. Once confirmed, we investigated environmental variables as potential determinants of the pattern. These climatic, seasonal and elevational variables were selected *a priori* based on our hypotheses.

For each of the 48 contiguous states, we determined mean daily, maximum, and minimum summer and winter temperatures; mean summer and winter precipitation; and average atmospheric pressure. To eliminate potential bias due to total rainfall, we also calculated the - difference between mean winter and summer precipitation (winter-summer). We obtained weather data using the Monthly Station Normals 1971-2000 CLIM 81 from NOAA and averaging across all weather stations within each state. Winter variables were calculated with data from December, January, and February, and summer variables used June, July, and August. Precipitation was used as a proxy for time of flowering: winter/early spring (cool season) vs. summer/early fall (warm season). Since changes in elevation also lead to changes in both temperature and atmospheric pressure, we used the barometric formula (eq. 1) with the annual mean temperature and the mean elevation of the state to determine mean atmospheric pressure (SI Table 1). Here, *P*_*h*_ is mean atmospheric pressure (in atmospheres), *P*_0_ is atmospheric pressure at sea level, *g* is the gravitational constant, *M* is molar mass, *T* is absolute temperature, *R* is the universal gas constant, and *h* is elevation in meters.

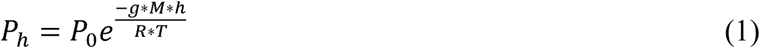

Mean elevation per state was taken from the 2004-2005 Statistical Abstract of the United States, Section 6.

General linear modelling was used to determine which variables correlated significantly with species richness. For each Family separately, we used a step-wise regression, eliminating at each step the least significant variables based upon their p-values. This left a linear model with the remaining significant variables at a level of p < 0.05. Because of redundancy among some variables, three different permutations of tests were performed. The first permutation used summer daily mean temperature, winter daily mean temperature, winter precipitation, summer precipitation, and atmospheric pressure. Since the west is drier with lower overall precipitation, we ran a second test using precipitation difference in lieu of winter and summer precipitation. A third test used daily maximum temperatures for hummingbirds and daily minimum temperatures for hawk-moths as the two Families are diurnal and nocturnal respectively.

#### Additional Species Richness Data

We can also provide figures for the species richness of hummingbirds by Canadian Provinces, Mexican States, and South American countries. We can provide species richness data for Canadian Provinces, six regions of Mexico (data by states is not available), Provinces of South Africa, and Australian Provinces.

#### Additional bird and insect Families of the continental USA

We specifically chose the hummingbird and hawk-moth Families as likely candidates for macrocompetition. Yet, what we find for each may simply be a more generally property of birds and insects in the 48 contiguous states, USA. So, we haphazardly selected 12 bird Families and 7 insect Families that exhibited sufficient species richness and data quality to test for within Family latitudinal and longitudinal trends (Table 3).

**Table 1:** GLM of Latitude and Longitude per state with species richness per family

| Family | *S* | Latitude | Longitude |
| --- | --- | --- | --- |
| Sphingidae | 101 | -1.301<sup>a</sup> | 0.264<sup>c</sup> |
| Trochilidae | 15 | -0.143<sup>a</sup> | -0.116<sup>a</sup> |

a: p<0.001, b: p<0.01, c: p<0.05

**Table 2:** Coefficients, total R-squared, and AIC for each final linear model. Permutation signifies the permutation of variables used for each regression as seen in the methods. PRECIP indicates precipitation, DIFF difference, SUM summer, WINT winter, AVG average daily mean temperature, MIN average daily minimum temperature, MAX average daily maximum temperature, and ATM atmospheric pressure

| Permutation | Family | Model | R<sup>2</sup> | AIC |
| --- | --- | --- | --- | --- |
| 1 | Sphingidae | $S = -7.701 + 0.116 * \text{SUM.PRECIP}^a + 1.470 * \text{SUM.AVG}^a$ | 0.6448 | 169.37 |
| | Trochilidae | $S = 8.215 - 0.021 * \text{WINT.PRECIP}^b - 0.052 * \text{SUM.PRECIP}^a + 0.249 * \text{WINT.AVG}^a$ | 0.651 | 47.64 |
| 2 | Sphingidae | $S = -20.73 - 0.094 * \text{PRECIP.DIFF}^a + 0.87 * \text{WINT.AVG}^a + 57.25 * \text{ATM}^a$ | 0.6187 | 173.76 |
| | Trochilidae | $S = 31.04 + 0.177 * \text{WINT.AVG}^a - 30.63 * \text{ATM}^a$ | 0.5149 | 62.96 |
| 3 | Sphingidae | $S = -5.90 + 1.89 * \text{SUM.MIN}^a$ | 0.6035 | 173.8 |
| | Trochilidae | $S = 28.54 + 0.164 * \text{WINT.MAX}^a - 28.98 * \text{ATM}^a$ | 0.5148 | 62.97 |

a: p<0.001, b: p<0.01

**Table 3:**
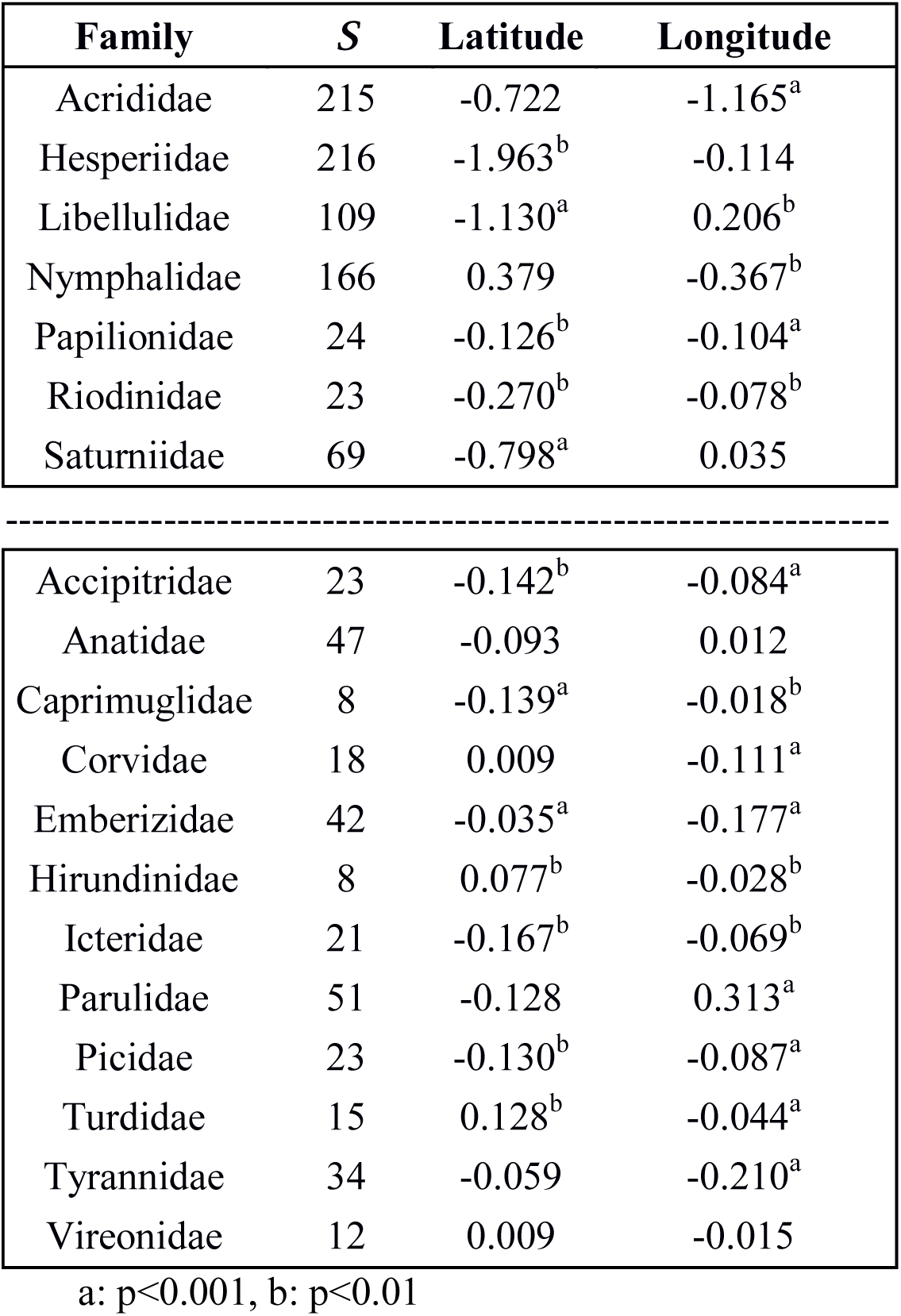
Linear model of Latitude and Longitude with species richness per insect and bird family

| Family | S | Latitude | Longitude |
| --- | --- | --- | --- |
| Acrididae | 215 | -0.722 | -1.165<sup>a</sup> |
| Hesperiidae | 216 | -1.963<sup>b</sup> | -0.114 |
| Libellulidae | 109 | -1.130<sup>a</sup> | 0.206<sup>b</sup> |
| Nymphalidae | 166 | 0.379 | -0.367<sup>b</sup> |
| Papilionidae | 24 | -0.126<sup>b</sup> | -0.104<sup>a</sup> |
| Riodinidae | 23 | -0.270<sup>b</sup> | -0.078<sup>b</sup> |
| Saturniidae | 69 | -0.798<sup>a</sup> | 0.035 |
| Accipitridae | 23 | -0.142<sup>b</sup> | -0.084<sup>a</sup> |
| Anatidae | 47 | -0.093 | 0.012 |
| Caprimuglidae | 8 | -0.139<sup>a</sup> | -0.018<sup>b</sup> |
| Corvidae | 18 | 0.009 | -0.111<sup>a</sup> |
| Emberizidae | 42 | -0.035<sup>a</sup> | -0.177<sup>a</sup> |
| Hirundinidae | 8 | 0.077<sup>b</sup> | -0.028<sup>b</sup> |
| Icteridae | 21 | -0.167<sup>b</sup> | -0.069<sup>b</sup> |
| Parulidae | 51 | -0.128 | 0.313<sup>a</sup> |
| Picidae | 23 | -0.130<sup>b</sup> | -0.087<sup>a</sup> |
| Turdidae | 15 | 0.128<sup>b</sup> | -0.044<sup>a</sup> |
| Tyrannidae | 34 | -0.059 | -0.210<sup>a</sup> |
| Vireonidae | 12 | 0.009 | -0.015 |

a: $p < 0.001$ , b: $p < 0.01$

## RESULTS

Fifteen hummingbird species and 101 hawk-moth species inhabit the continental United States. Figures 2a and 2b show the species richness of hawk-moths and hummingbirds, respectively, in the United States, Canada, and Mexico by state, province, territory, and region. Inspection of these graphs reveals that hummingbird species increase from north to south and from east to west (Figure 2b). Hawk-moth species likewise increase from north to south, but in contrast to hummingbirds, hawk-moth diversity increases from west to east (Figure 2a). The result of the latitudinal and longitudinal GLM confirm the directional bias seen on the map (Table 1). In summary, hummingbird diversity peaks in the Southwestern United States, while hawk-moth richness peaks in the Southeastern United States.

**Fig. 1.**
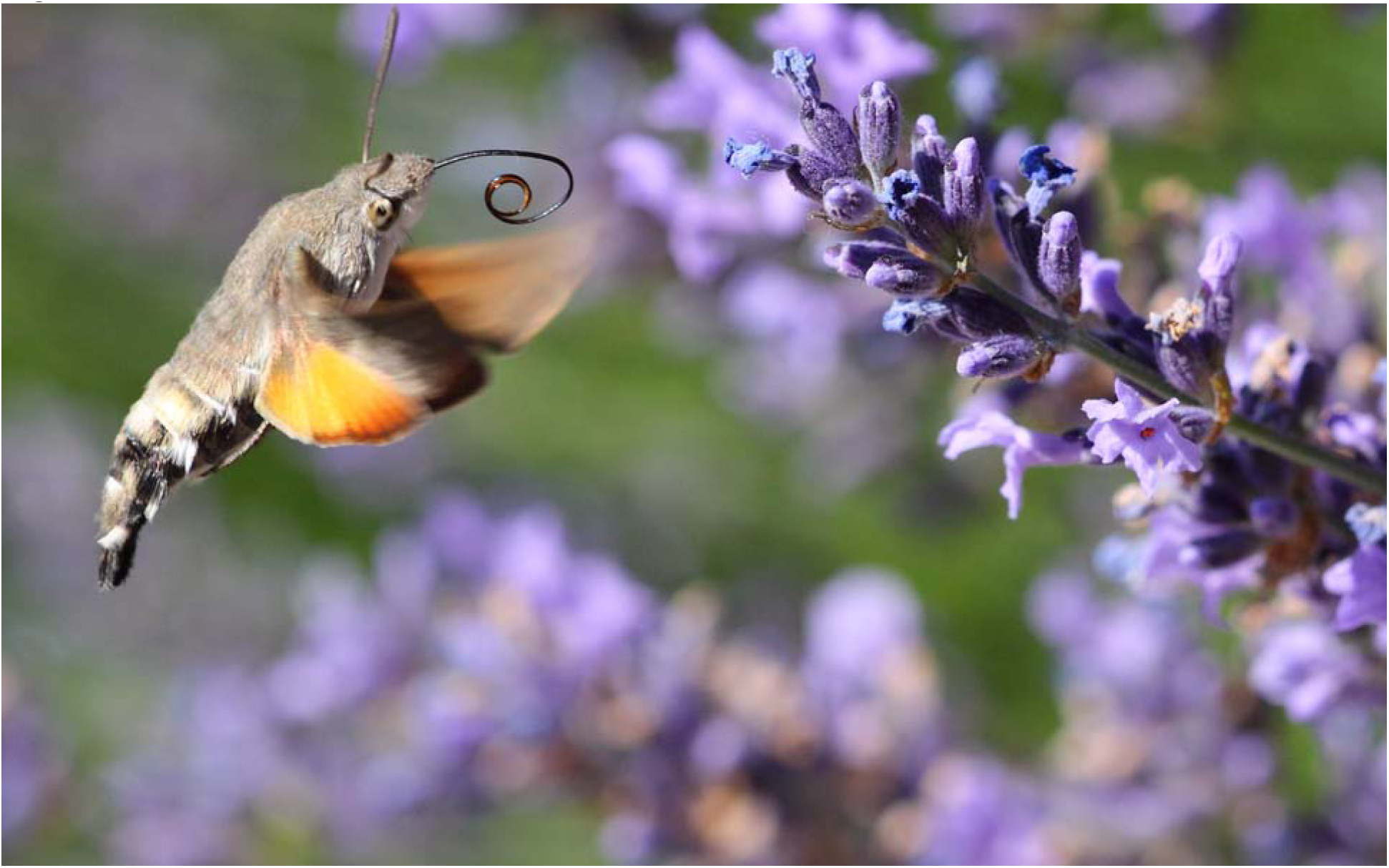
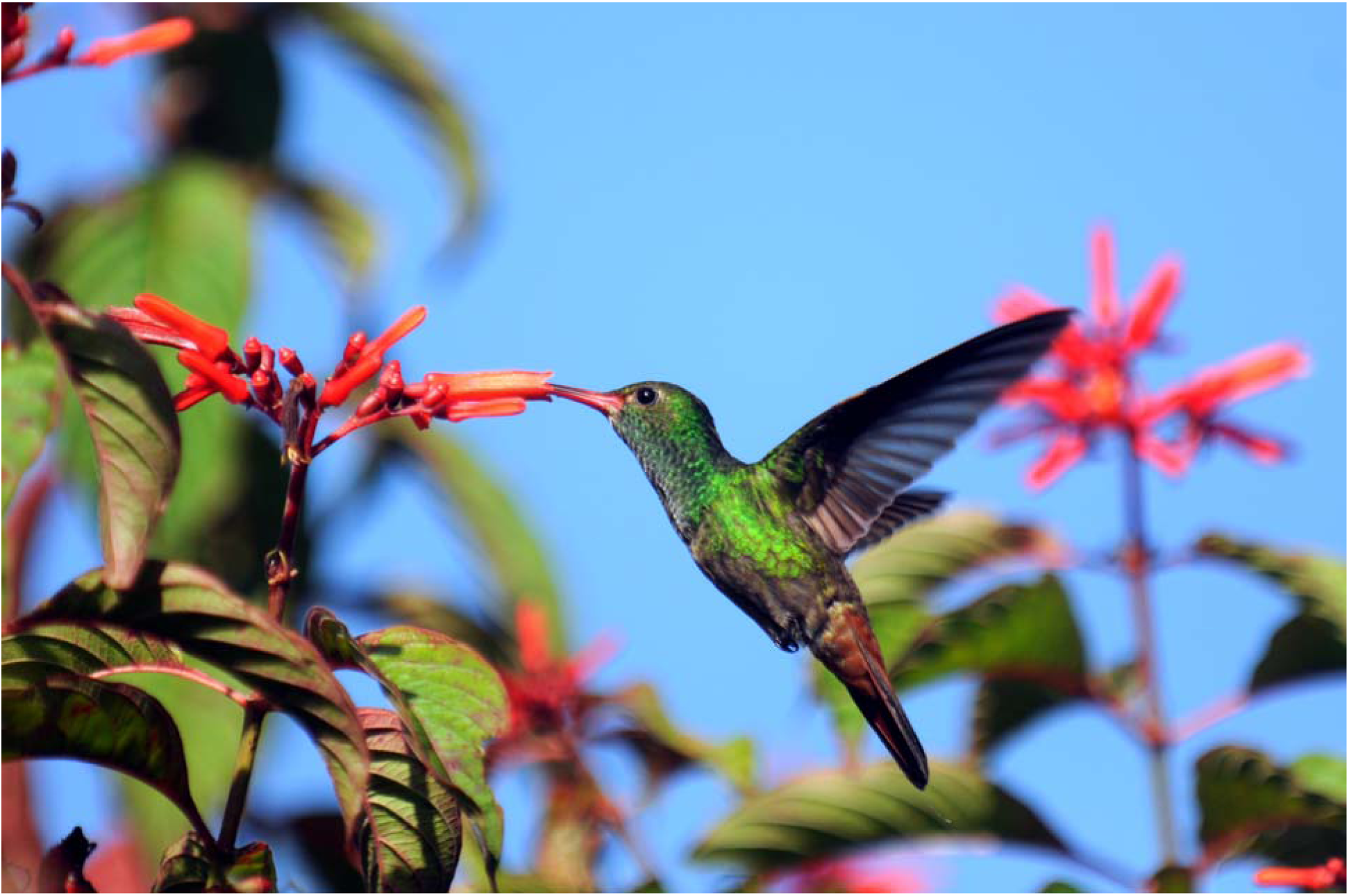
(a) *Macroglossum stellatarum*, the Hummingbird Hawk-moth, hovering by lavender flowers (b) *Amazilia tzacatl*, the Rufous-tailed Hummingbird, feeding in Costa Rica. As seen in the image below, hawk-moths have extremely long extensile probosces for collecting nectar and the ability to hover in front of flowers. Convergent features are seen in the image of the hummingbird below with long bills and extensile tongues and the ability to hover. Image of *M. stellatarum* by Thorsten Denhard, CC-BY-SA-3.0. Image of *A. tzacatl* by T. R. Shankar Rama, CC-BY-SA-4.0

**Fig. 2.**
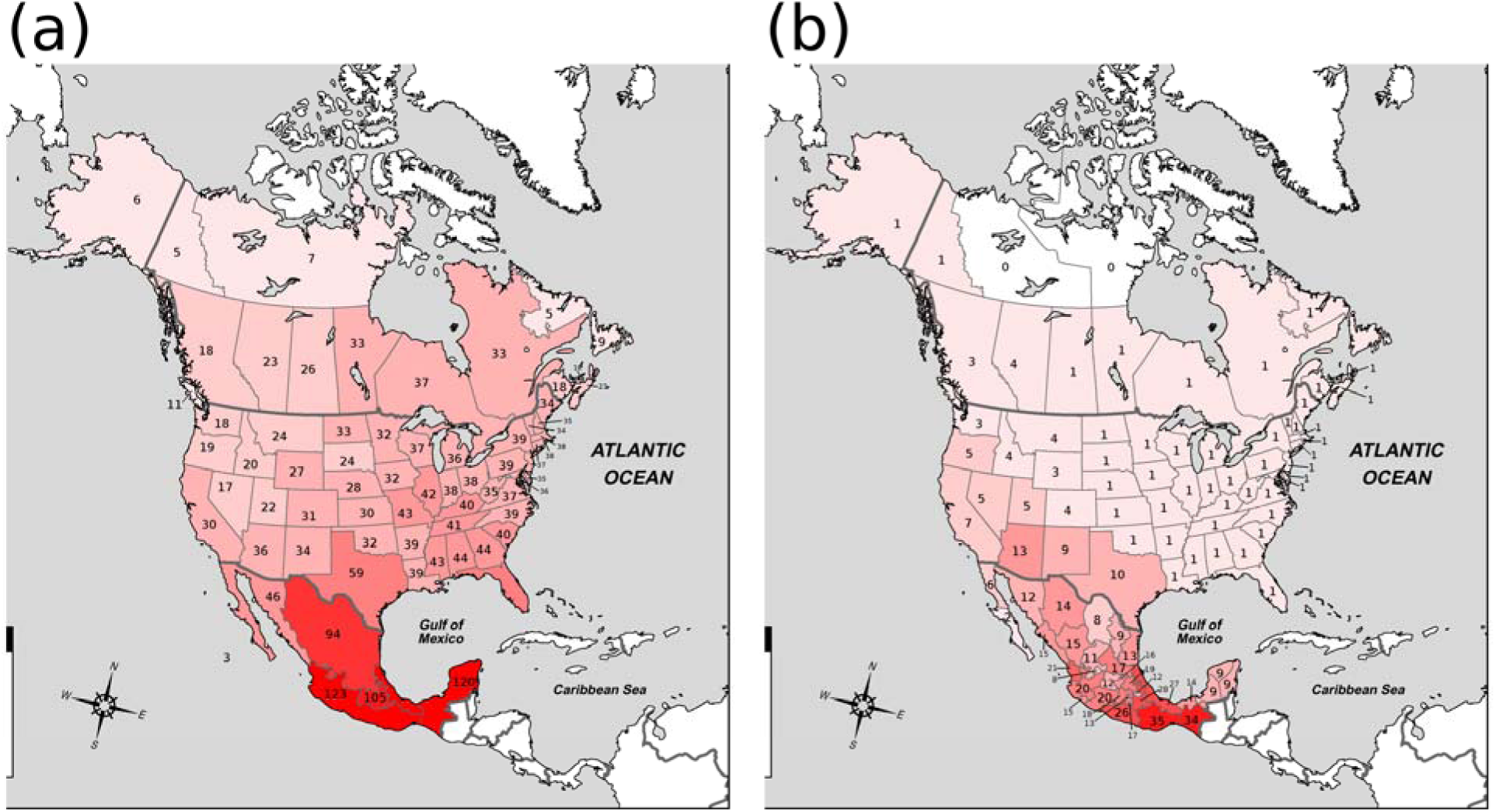
Species richness per state, province, territory, and region of in the United States, Canada, and Mexico (a) hawk-moths (order Lepidoptera, family Sphingidae) and (b) hummingbirds (order Apodiformes, family Trochilidae). Greater color intensity reflects greater proportional species richness per family. As one can see, hawk-moths are more species rich in the eastern half of the northern North American continent while hummingbirds are more species rich in the western half of the northern North American continent. Both species show increasing species richness moving from north to south.

We examined the association between environmental variables and the diversities of hawk-moths and hummingbirds with the results shown in Table 2. Model permutation 1 shows hawk-moth species richness increasing with summer precipitation and summer daily average temperatures, and hummingbird species richness decreasing with winter and summer precipitation and increases with winter daily average temperature. With permutation 2, the relationship for hawkmoths now showed an increase with winter daily average temperature and atmospheric pressure and a decrease with precipitation difference, indicating an association with summer rains; hummingbird species richness showed a positive correlation with winter daily average temperature and a negative correlation with atmospheric pressure. The third permutation of tests, which used the minimum and maximum temperatures for hawkmoths and hummingbirds respectively, showed similar correlations for hummingbirds as the second test (but with new coefficients) but that hawk-moths were only significantly correlated with summer daily minimum temperature. In summary, hawk-moth diversity is higher in states with higher summertime precipitation and temperatures, particularly the minimum temperature (consistent with nocturnal activity), and states with overall low elevations. Hummingbird diversity is higher in states with higher wintertime temperatures, particularly the winter highs (consistent with diurnal activity), and states with higher elevation.

### Additional Species Richness Data

The latitudinal trends in species richness of hawk-moths and hummingbirds is pervasive throughout North America, from Canada to Mexico (Fig. 2). Only one species of hummingbird is found east of the province of Alberta, Canada, while hawk-moth species richness is greater in eastern than in western Canada. The small eastern province of Prince Edward Island, for instance, has the same number of species as the large western province of British Columbia. The geographical trend is more difficult to see in Mexico, because hawk-moth species richness could only be resolved to 6 regions. Nonetheless, hummingbird species richness in the Yucatan is low compared to close-by and neighboring states to the west. Each state (Yucatan, Campeche, and Quintana Roo) has only 9 species each – 10 combined – compared with 14 in neighboring Tabasco, 20 in Michoacán, and 26 in Guerrero. In contrast, there are 133 hawk-moth species present in the state of Veracruz alone compared to 120 species in the states of Nayarit, Jalisco, Colima, Michoacán, Oaxaca, and Guerrero combined.

Species richness of hawk-moths and hummingbirds appears to exhibit a similar reversal in directional dominance in South America as we found in North America. Hummingbird species richness is greater in western than in eastern South America (Johnsgard, 1997; NatureServe, 2010). Ecuador has approximately twice as many species as Brazil, and hummingbird species richness per area is greater in Chile than in Argentina despite having fewer species overall.

Though the data for hawk-moths in South America are less comprehensive, species richness appears greater in French Guiana, Argentina, Bolivia, and Venezuela (CATE, 2010) than in countries to the west. Together with the results from North America, climate and topography appear to exert opposing effects on the continental distributions of hawk-moths and hummingbirds. Finally, both South Africa and Australia (Fig. 3) show support for hawk-moth species richness increasing longitudinally from west to east, and latitudinally towards the equator. What we do not known is whether these north-south and east-west patterns hold up within the very large Cape and Gauteng Provinces of South Africa and the Western Province of Australia.

**Fig. 3.**
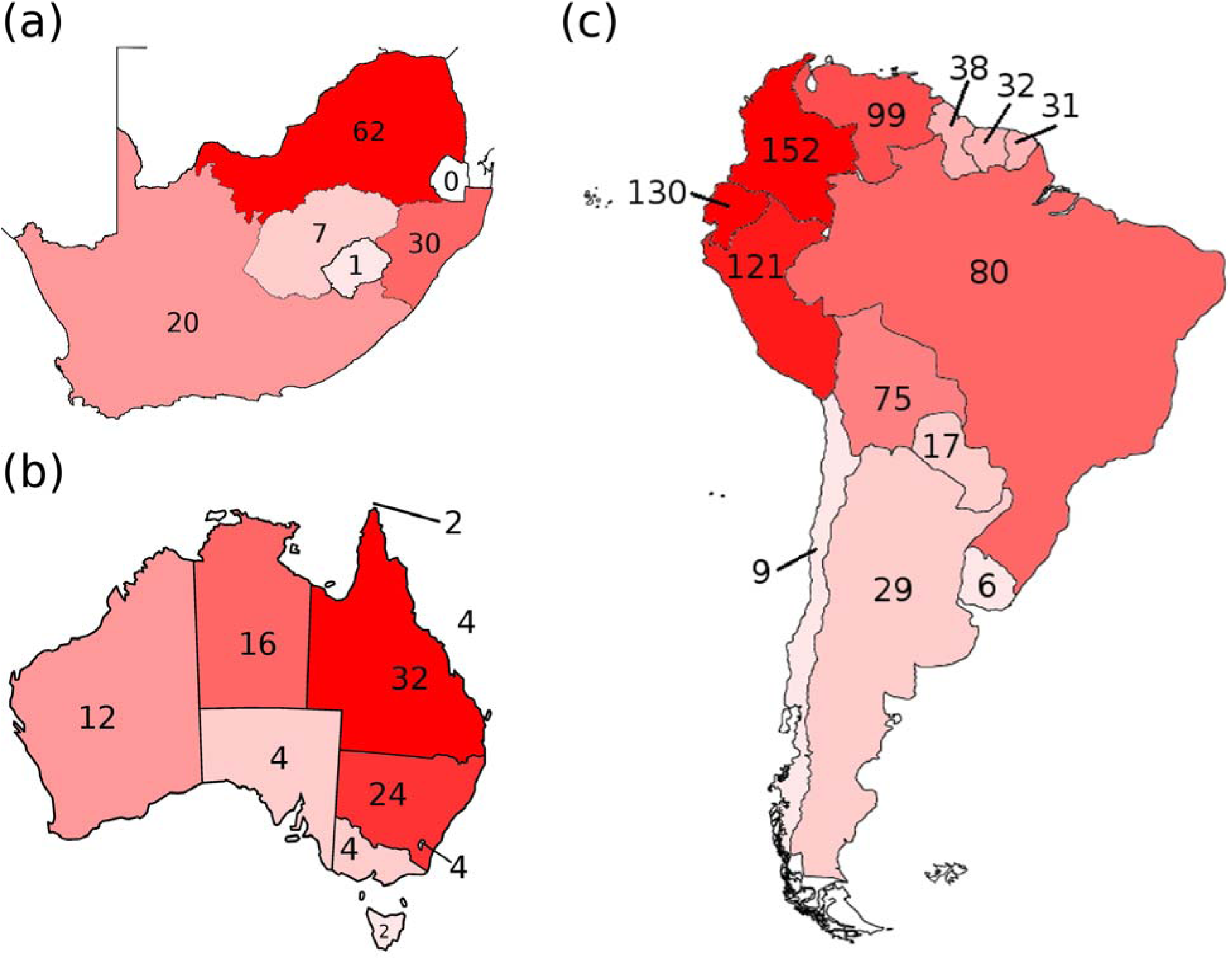
Species richness of a) hawk-moths per pre-1994 South Africa province and b) hawk-moths per state and territory of Australia, and c) hummingbirds per country in South America. One can clearly see the same eastern bias for hawk-moths and western bias for hummingbirds.

### Additional bird and insect Families of the continental USA

Of the seven additional insect Families two showed no significant latitudinal trends, four showed significantly increasing richness from east to west (opposite hawk-moths), and only one shared the same pattern as the hawk-moths (Table 3). The species richness of Libellulidae (largest family of dragonfly) increases from west to east. Of the twelve bird Families, two showed no significant latitudinal patterns, one showed a significant increase in richness from west to east (the Parulidae), and nine exhibit the same pattern as the hummingbirds with more species as one moves westwards.

Despite shared trends the hummingbirds do stand out in having just a single species in 36 of 48 states of the contiguous USA. All of the other bird families of Table 2 have at least three or four species reaching east coast states, even those with fewer total numbers of species than hummingbirds.

## DISCUSSION

Our analyses for evaluating evidence for macrocompetition between hummingbirds and hawk-moths has its limitations. Firstly, correlation does not necessarily illuminate causation. Yet, given the degree to which hawk-moths and hummingbirds have been ignored as potential shapers of each other’s biodiversity and distributions, correlations will shed light on some of our hypotheses and suggest new ones. Secondly, using states of the USA as our unit of replication is geographically crude; they vary in size by more than two orders of magnitude, have diverse and irregular shapes, and adjacent states will have some degree of spatial autocorrelation. A more fine-grained and detailed level of division, such as the county, or the use of GIS data would be preferable, and in many cases possible for hummingbirds. Unfortunately, hawk-moth diversity and distribution data are coarse; and this is the first systematic analysis of hawk-moth species richness. Fine grain data on hawk-moth species’ ranges and presence/absence are deficient throughout the world and digital range maps are scarce to non-existent with the range maps from Tuttle (2007) not digitizable. We feel our scale best balances the need for accurate data and diverse sampling units. We have high confidence in state-wide inventories of hawk-moths. Perhaps, intriguing results will inspire more detailed interest and work. Thirdly, due to the irregularities of species boundaries, states, especially those along the border with Mexico, could contain species with well-established populations that occupy just a fraction of the state. While this inflates the numbers of species within the state, any boundary drawing would necessarily have this problem. We felt it best to accept the current haphazard sizes and irregularities of states rather than create more regular spatial sampling schemes that would amplify uncertainties in the presence/absence of hawk-moth species.

Despite these limitations, our results show clearly divergent trends in broad scale geographical patterns of hawk-moths and hummingbirds and are striking and consistent with the hypothesis of intertaxon competition and a phenomenon we call macrocompetition. Environmental correlates of the geographic trends for each of the two taxa further support the potential for geographical partitioning based on elevation, precipitation, and temperature. Though only suggestive, our investigation highlights an important yet understudied interaction at higher levels of taxonomy. The pattern for hummingbirds is well known, but the pattern that emerged for hawk-moths is novel and quite unexpected.

### Basic Geography

The 15 hummingbird species and 101 hawk-moth species in the US exhibit a clear and opposite latitudinal bias in species richness. Most hummingbird species are found in the western United States, with only 1 species breeding east of the Rocky Mountains. This result is generally consistent with the expectation that species diversity should be greater in mountainous areas due to habitat heterogeneity and reproductive isolation. The distribution of hawk-moths on the other hand offers some striking and initially counter-intuitive patterns. Despite their relatively large size and habitat heterogeneity, western states are conspicuously depauperate in hawk-moth species. States on the eastern seaboard just a fraction of the size of California have much higher hawk-moth diversities. Similar diversity asymmetries appear when noting Maine’s (extreme northeast) high diversity in contrast to low diversity in the state of Washington (far northwest). Even neighboring states show this pattern with North Dakota having over 25% more hawk-moth species than Montana despite being smaller in area, sharing a biome, and offering much less environmental heterogeneity. This clear portioning invites explanation by way of climatic analysis.

### Climate Analysis

Being a large, nocturnal insect poses challenges and opportunities for such an evolutionary technology. Accordingly, the analyses showed how hawk-moth species richness increases with higher temperatures – especially high summer minimums – higher summer precipitation (associated with warm season flowering), and high atmospheric pressure. In studies along mountain gradient, hawk-moth pollination activity fell off once summer minimums reached below 15°C in mountainous areas (Harlington, 1968; Cruden et al., 1976). With respect to elevation, insects likely suffer more than birds from a drop in the partial pressure of oxygen; the hawk-moth tracheal system requires diffusion of O_2_ into and CO_2_ out of spiracles located on the exoskeleton. Adult insects, as shown by a study with the tobacco hornworm, *Manduca sexta*, are smaller when reared under hypoxic conditions (Harrison et al., 2010). And, smaller size is not favorable for hawk-moths that must maintain thoracic heat for flight (Dorsett, 1962). As seen in North America, South Africa, Australia, and likely South America, the eastern sides of continents are more likely to offer summer rains and higher nighttime temperatures during the summer relative to west sides.

Relative to hawk moths, the evolutionary technology of hummingbirds should allow for a higher fitness in colder temperatures, cool season flowering, and lower barometric pressures both in regard to temperatures and oxygen partial pressures. Species richness of hummingbirds correlated with all of these in the predicted direction. In the Andes of South America, hummingbird richness is highest in the 1800-2500m range (Schuchmann, 1999) as well as highest between 500-1800m in southwestern United States (Wethington et al., 2005). The respiratory system of hummingbirds maximizes the intake of O_2_. Correcting for body size, hummingbirds have the largest heart, fastest heart and breathing rates, and densest erythrocyte concentrations among all bird families (Johnsgard, 1997). For example, *Colibri coruscas* increases oxygen consumption by only 6 to 8% when hovering under hypoxic conditions equivalent to an altitude of 6000m (Berger, 1974). Hummingbirds are better pollinators than insects in the cloudy, windy, and rainy conditions often found at high elevations (Cruden, 1972). The species richness of hummingbirds at these altitudes may result from their physiological adaptations (McGuire, 2014). For somewhat similar reasons of inter-taxonomic, ant diversity may steadily decline with elevation while in the tropics small mammal diversity often peaks at intermediate elevations (Nor 2001, Heaney 2001, McCain 2005, Sanders et al. 2007).

Different evolutionary technologies should influence methods of resource exploitation as well as the costs. We can expand the notion of fundamental and realized niches to whole Families. It is clear that hawk-moth characteristics favor areas of warm growing season flowering that are low in elevation and oxygen rich. Presumably these are features that would favor hummingbirds as well. Though endotherms, hummingbirds are extremely small and easily lose heat. Many hummingbird species undergo nightly torpor to conserve energy. Moreover, though suffering only a small uptick in energetic costs at higher elevations, hummingbirds are still more efficient at lower elevations. Furthermore, migratory hummingbird species, like hawk moths, time their activity in the United States for the summer. It seems that the fundamental niche of hummingbirds contains that of hawk-moths, and extends further into colder temperatures and higher elevations. So, why does the species richness and hummingbird Family’s realized niche not match its fundamental niche? *Why only one hummingbird species east of the Rocky Mountains*?

### Whither Macrocompetition?

Hummingbirds and hawk-moths are two phylogenetically distant, yet morphologically convergent families of nectarivores (Cruden et al., 1983; Carpenter, 1979; Kessler et al., 2010; Aigner and Scott, 2002; Fulton, 1999; Willmott and Burquez, 1996). Prior work supports high niche overlap and definitive micro-competition and the opportunity for meso-competition. Our analyses support geographical partitioning and inter-taxonomic macrocompetition. That hawk-moths contribute to an unusually low diversity of hummingbirds in the east and vice-versa in the west provides an open but intriguingly viable hypothesis.

The evolution and distribution of nectar feeding bats in the family Phyllostomidae may also shed light on the possibility of macrocompetition between hummingbirds and hawk-moths. These bats use the same class of resources, taking from flowers with similar – if not higher – amounts of sugar and nectar to those pollinated by hawk-moths and hummingbirds (Cruden et al., 1983). The biogeography of these bats resembles that of hummingbirds, existing primarily in tropical regions (none inhabit temperate North America) with species richness highest in the Andes. Furthermore, Phyllostomid bats underwent a radiation in the mid-Miocene around the same time as the hummingbird radiation. Though diversifying in a similar manner to hummingbirds, there are many fewer species of true nectarivores — sixteen genera with 38 species adapted for nectar feeding (Fleming et al., 2009). This relatively limited species diversity of bat nectarivores may reflect competition from both hummingbirds and hawk-moths for available niche space. These nectarivorous Phyllostomid bats show a similar pollination syndrome to hawk-moths – nocturnal foraging with dependence on scent to guide them to flowers – with some plant species relying on both bats and hawk-moths for pollination (Hernandez-Montero and Sosa, 2015). We conjecture that bat nectarivore diversity is also constrained by competition from hawk-moths at lower elevations and by hummingbirds at higher elevations, severely restricting their potential niche space to high-elevation areas at night.

The alterative to macrocompetition would view the geographic partitioning of hummingbirds and hawk-moths as apparent and not causally related. Indeed, a number of North American bird Families show east to west increases in diversity, and perhaps this is the norm. In particular, we do not suggest that showing geographic partitioning by haphazard pairs of Families provides evidence for macrocompetition. Here the two Families were selected *a priori*, because of their convergence in morphology and ecological niche similarities. They do exhibit microcompetition (Carpenter 1979). In terms of mesocompetition, experiments are lacking as to whether within communities hawk-moths depress the population sizes of hummingbirds and vice-versa. But, given the capacity of hawk-moths and hummingbirds to deplete nectar, it seems likely. The geographic partitioning of diversity was striking. No bird Families with comparable total continental diversities plunge to just one species as one hits the longitudinal midsection of the United States; and the results for the hawk moths are novel and striking in the opposite direction. Of seven other insect Families only one exhibited a significant increase in species richness from west to east.

In terms of Brown and Davidson’s (1979) requirements for inter-taxonomic competition the first and third indicators are met. While not conclusive, the evidence for macrocompetition between hawk-moths and hummingbirds is tantalizing. The all but ignored eco-evolutionary relationships between hawk-moths and hummingbirds seems ripe for study. Profitable avenues of studies could include 1) better local data on hummingbird and hawk-moth diversities and population sizes, and 2) mesocompetition studies on smaller scales to verify that each Family can influence local abundances and diversities. Looking more generally, we hope our analyses inspire further study into local and even continent-wide distributions of competition and niche partitioning.

## ACKNOWLEDGEMENTS

Abdel Halloway wishes to thank the NSF for funding his graduate studies. This material is based upon work supported by the National Science Foundation Graduate Research Fellowship under Grant Nos. DGE-0907994 and DGE-1444315. Any opinion, findings, and conclusions or recommendations expressed in this material are those of the authors(s) and do not necessarily reflect the views of the National Science Foundation.

